# CGMFinder Identifies Correlated Gene Modules from 3H scRNA-seq Data

**DOI:** 10.1101/2024.11.29.624642

**Authors:** Wenping Ma, Xiaoliang Sunney Xie

**Affiliations:** Changping Laboratory; Beijing, 102206, China; Beijing Advanced Innovation Center for Genomics (ICG) & Biomedical Pioneering Innovation Center (BIOPIC), Peking University; Beijing 100871, China

## Abstract

Correlated gene modules (CGMs) contain genes whose expression fluctuates together. Genes in CGMs are often functionally related and regulated by shared transcription factors. CGMs can be identified under steady-state conditions in populations of cells using single-cell RNA sequencing (scRNA-seq). Here, we introduce CGMFinder, a tool for CGM identification using “3H” scRNA-seq data (High mRNA capture efficiency, High cell numbers, and High sequencing depth). CGMFinder employs a graph-based filtering approach, first identifying CGM cores from highly-expressed genes and then linking noisy low-abundance genes to these cores. In lymphoblastoid cell line 3H datasets generated by in-lab and commercial protocols, CGMFinder accurately identifies CGMs enriched for gene ontologies or pathways. In cells grown under hypoxic conditions, CGMFinder successfully identified hypoxia-specific “glycolysis” and “response to oxygen levels” modules. Evaluations using ground truth correlation modules demonstrate that CGMFinder outperforms other CGM identification methods such as WGCNA and FastICA in scRNA-seq data.

## Main

Genes do not operate in isolation. Instead, they frequently work together to form modules that carry out specific biological functions. Identifying and characterizing these modules is important for gaining an understanding of biological systems^1,2^. High-throughput gene expression data, such as microarray and bulk RNA sequencing, have commonly been used to identify CGMs based on similar expression patterns^3,4^. However, these population-level data represent average gene-gene relationships, often across diverse cell types mixed in the sample. Single-cell RNA-seq (scRNA-seq) technologies^5,6^ have transformed transcriptome profiling by enabling the analysis of thousands of individual cells in a single experiment. This allows researchers to identify CGMs for a specific cell type using single-cell data (Fig. 1a). Despite these exciting opportunities, the use of scRNA-seq also introduces several challenges, which add complexity to the identification of CGMs.

**Fig. 1.**
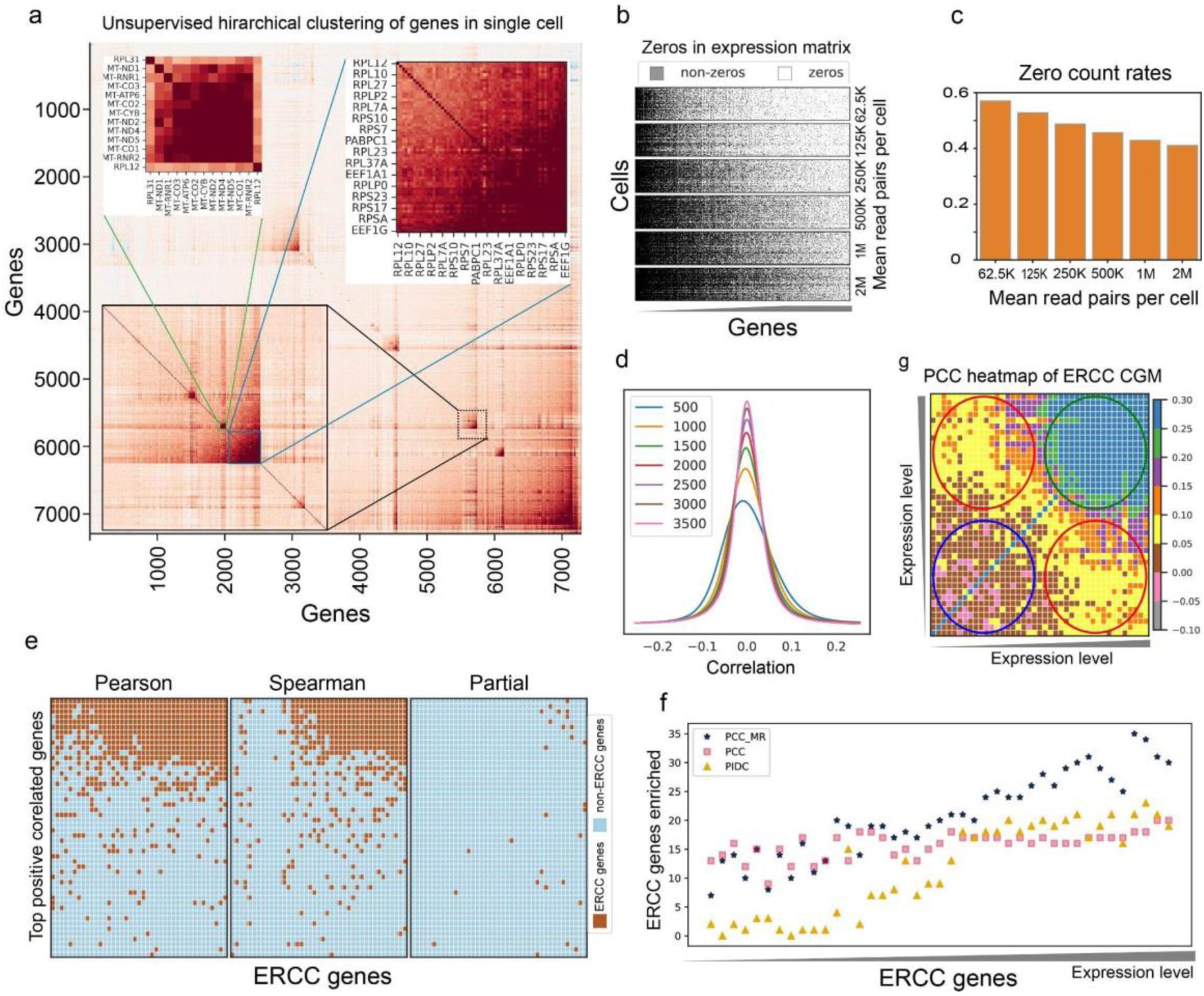
Selection of ground-truth modules and gene correlation measurement. **a,** Illustration of correlated gene clustering using scRNA-seq data from 500 GM12878 cells by unsupervised hierarchical clustering. Genes exhibiting highly similar expression across individual cells are clustered into modules. The inset plots show two such clusters comprising functionally related genes: ribosomal protein genes (blue box) and mitochondrial-encoded genes (green box). **b,** Zeros in the gene expression matrix at varying sequencing depths sampled from the 500-cell GM12878 data. **c,** Zero count rates across different sequencing depths from the 500-cell GM12878 data, with genes expressed in fewer than 10% of cells removed. **d,** Pairwise PCC distribution across a varying number of cells sampled from the GM12878 dataset at 2M sequencing depth.. **e,** Top 50 neighbors for the ERCC module using PCC, SPC, and partial correlation respectively. Brown dots indicate ERCC genes and light blue dots represent non-ERCC genes. **f,** Enrichment of ERCC genes in the top 50 positive correlated neighbors for each ERCC gene using Pearson’s correlation coefficient (PCC), Mutual Ranking of PCC(PCC-MR), and Partial Information Decomposition (PIDC) as measurements. ERCC genes are sorted by increasing expression levels. **g,** Expression level-associated correlation bias for the ERCC module. ERCC genes are sorted by ascending expression levels, and PCC was used for correlation measurement. The green circle indicates the highly correlated “module core” from highly-expressed genes, the red circles indicate moderate positive correlations between highly expressed genes and lowly expressed genes, and the blue circle indicates weak positive or negative correlations between lowly expressed genes, as described in the main text.

One of the primary challenges in scRNA-seq data analysis is dealing with sparse data^7^. Sparse data is caused by factors such as low transcript abundance and the inaccuracy of current methods of transcript quantification. The process of identifying CGMs from scRNA-seq data depends initially on measuring pairwise correlations between genes^8^. The accuracy of module detection is highly dependent on the precision of these pairwise gene correlation measurements. Data sparsity directly impacts the accuracy of these correlations, with expression-level dependent correlation biases being of particular concern.^9,10^.

Protocols with high mRNA capture efficiency and increased sequencing depth have been developed to enhance transcript detection, especially for low-expression genes, thereby reducing the impact of sparsity on pairwise correlation measurements^11–14^ ^15^ (Fig. 1b and Fig. 1c). Another critical factor influencing the accuracy of pairwise correlation calculations in scRNA-seq data is the number of profiled cells. A larger cell sample size provides greater statistical power for correlation assessments^16^. With a limited number of cells, uncorrelated genes may mistakenly be identified as correlated^15^ (Fig. 1d).

Several methods have been developed for constructing co-expressed gene networks, including WGCNA^19^, GENIE3^20^, GRNBOOST2^21^, PIDC^22^, SCENIC^23^, and scLink^24^. After constructing co-expressed networks, CGM identification is typically performed using approaches such as clustering, decomposition, and biclustering^29^. However, benchmarking studies assessing the performance of network-building tools on single-cell data suggest that there is room for improvement ^25–28^, and tools that can seamlessly integrate network construction and module identification are still limited.

Here, we developed a novel method called CGMFinder (**C**orrelated **G**ene **M**odule identi**Fi**cation by expressio**n**-weighte**d** int**er**connection) for more accurate identification of CGMs within scRNA-seq datasets. Accurate detection of CGMs relies on high-precision 3H data from a uniform population of cells, so we employed MALBAC-DT^15,30^ (Multiple Annealing and Looping Based Amplification Cycles for Digital Transcriptomics) a high-performance in-house protocol to generate a 3H dataset for the homogeneous GM12878 cell line. To create standards for objective evaluation of module detection and to optimize correlation measurement selection, we employed two “ground truth” modules, consisting of external spike-in standards and internal mitochondrial operon-like genes. We used this 3H dataset and the evaluation standards to develop and fine-tune CGMFinder. CGMFinder implements a graph-based approach that identifies CGMs by assuming that genes within a module are tightly connected, allowing noisy inter- and intra-module connections to be removed. This method requires only the gene expression matrix as input and enables accurate assignment of lowly-expressed genes to modules. CGMFinder also includes a method for calculating the distance between modules, facilitating module-level analysis of functional structures within scRNA-seq data. Using the latest commercial single-cell platform (10x genomics v4), we generated another 3H dataset for the same GM12878 cell line with a larger number of cells. CGMFinder detected a higher number of modules in this dataset, and the modules were highly similar to those identified in the MALBAC-DT dataset. We produced an additional 3H dataset for a GM12878 sample under hypoxic stress using the 10x platform. CGMFinder successfully identified the hypoxic-specific “glycolysis” and “response to oxygen levels” modules in this dataset, which were not detected in the normoxia dataset. Finally, we compared CGMFinder with other module detection methods by evaluating the identification of the two ground-truth modules, demonstrating the superior performance of CGMFinder over other methods.

## Results

### Selection of ground-truth modules and gene correlation measurement

The selection of gene correlation measurements serves as a vital initial step in network construction. Numerous correlation measures have been proposed to evaluate the pairwise relationships among genes, including widely recognized techniques such as Pearson’s correlation coefficient (PCC), Spearman’s correlation coefficient (SCC), partial correlation, and PIDC’s partial information decomposition^31^. Ranked correlation methods such as Mutual Ranking (MR) and Highest Reciprocal Ranking (HRR) have also been employed, and have shown good performance in gene co-expression analysis^32,33^. Despite numerous literature reviews offering recommendations for different association measures^34–39^, the selection of an appropriate measure remains challenging due to the lack of standardized evaluation approaches.

To address this challenge, we introduce two ground-truth modules: the “external ERCC module” and the “internal mitochondria module” to assess the selection of association measures and evaluate the precision of CGM identification. For an external standard, we added 92 ERCC (Evaluation of the External RNA Controls Consortium) RNA spike-ins to the cell lysis mix, which was robotically distributed to each well of a 96-well plate (Methods). The “co-expression” characteristic of the ERCC external molecules arises due to their consistent ratio to each other and the uniform quantity of each RNA introduced into every cell. The 92 ERCC spike-ins used in this study are strategically designed to cover a broad spectrum of transcript abundance levels, making them an ideal tool for evaluating the variability present in genes with varying expression levels^40^. For an internal standard, we used the co-regulation of genes encoded by the mitochondrial genome, which is based on their transcription from lengthy polycistrons under the control of distinct promoters^41^.

We hypothesize that effective correlation measures should position module members prominently among their top-ranked correlated neighbors. We conducted an initial evaluation of the efficacy of three correlation measures (PCC, SCC, and partial correlation) using these two ground-truth modules. We ranked correlation values for each gene and assessed the enrichment of module members within the top fifty nearest neighbors. Our analysis revealed that both PCC and SCC outperformed partial correlation in both the ERCC module and mitochondria module (Fig. 1e and Extended Data Fig. 1a). Particularly, PCC displayed superior performance to SCC in the ERCC module for genes with lower expression levels (Fig. 1e). Conversely, the performance of PCC and SCC for the mitochondria module was comparable, given that all genes within this module exhibit high expression levels (Extended Data Fig. 1a).

Subsequently, we compared the performance of PCC with two other methods (PCC-MR and PIDC) using the ERCC module. In general, PCC-MR demonstrated superior performance for highly expressed genes among these three approaches, while showing similar results to PCC for genes with lower expression levels. PIDC yielded better outcomes for highly expressed genes than PCC, but displayed the worst performance for genes with lower expression levels (Fig. 1f). Within the mitochondria module, these three methods produced results similar to those in the ERCC module (Extended Data Fig. 1b). Based on these comparative analyses, we selected PCC-MR as the correlation measure for this study.

### Principle and workflow of CGMFinder

It is important to note that correlation measurements showed diminished performance when applied to genes with low expression levels. To explore how expression levels influence noise, we analyzed the PCC heat map of genes within the ERCC module. Our null assumption was that, in the absence of noise, correlations among ERCC spike-in RNAs would remain consistently positive. However, the analysis revealed three distinct correlation patterns: (Ⅰ) Highly expressed genes exhibited strong positive correlations with one another, forming a tightly interconnected “module core” (Fig. 1g, green circle). (Ⅱ)Genes with low expression levels demonstrated weak positive or even negative correlations within their group (Fig. 1g, blue circle). (Ⅲ) Moderate positive correlations were observed between highly expressed genes and lowly expressed genes (Fig. 1g, red circle). This suggests that correlations among lowly expressed genes within the same module may be obscured, but their membership in the module can still be inferred through connections with highly expressed genes.

Managing noisy correlations from lowly-expressed genes is crucial for precise module detection. Neglecting this issue can lead to incorrect gene assignments and the generation of false positives. Based on the observation of biased correlation patterns in the ERCC module, we developed CGMFinder to detect CGMs in scRNA-seq data. To address the noise associated with lowly expressed genes, CGMFinder identifies modules by leveraging strong connections among highly expressed genes to define module cores and then assigns other genes to modules based on their connections with the existing members.

The CGMFinder workflow integrates co-expressed gene network construction and module detection into a seamless process, requiring only the gene expression matrix as input (Fig. 2a). First, the approach computes the PCC matrix from the expression data (Fig. 2b) and converts it into a PCC-MR (PCC-Mutual Ranking) matrix, which acts as a distance matrix (Fig. 2c). Rather than applying hard or soft thresholding, CGMFinder selects the top N most correlated genes from the PCC-MR matrix for each gene, ensuring the equal inclusion of lowly expressed genes in the network (Fig. 2d). The network edges are sorted by ascending MR values and grouped into bins to enable sequential use of edges from low to high MR values during module detection. (Fig. 2e). To identify modules, edges within each bin are progressively added to construct the “raw network” (Fig. 2f, bottom layers; Fig. 2g, left). To eliminate unstable or noisy connections, this network is then refined by requiring that each edge is the side of a triangle, resulting in the “triangle network” (Fig. 2f, middle layers; Fig. 2g, middle). The triangle network is then further refined by the removal of non-consecutive nodes, which are defined as those whose removal causes clusters to split into at least two parts (Extended Data Fig. 2). Removal of these genes creates the final “consecutive network” (Fig. 2f, top layers; Fig. 2g, right), where new modules are identified.

**Fig. 2.**
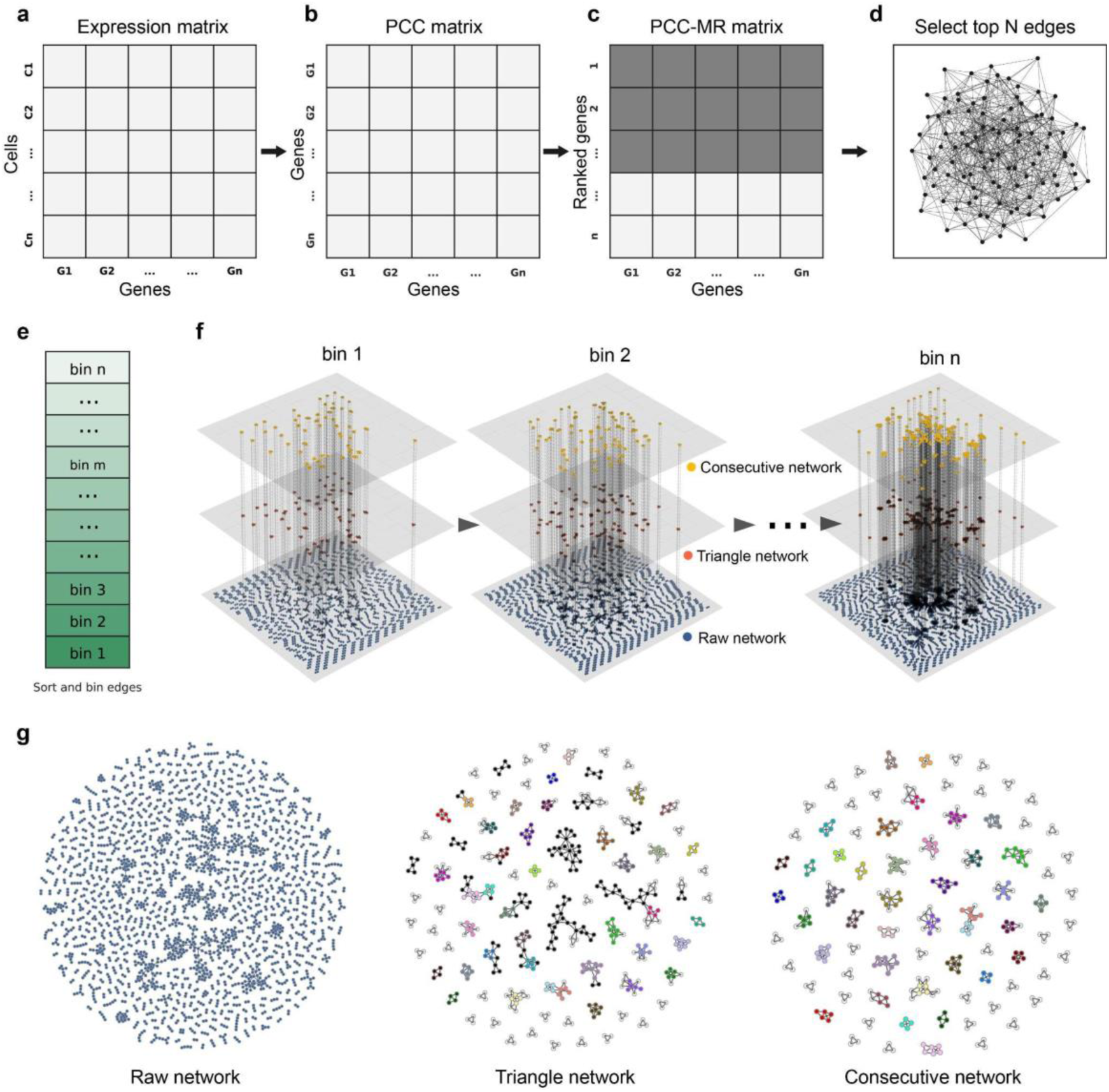
Schematics of principle and workflow of CGMFinder. **a,** Generation of the gene expression matrix from scRNA-seq data. **b,** Calculation of the pairwise Pearson Correlation Coefficient (PCC) matrix from the gene expression matrix. **c,** Calculation of the PCC-Mutual Ranking (PCC-MR) matrix from the PCC matrix. Shaded boxes represent the top neighbors. **d,** Construction of the co-expression network using edges from the top N neighbors for each gene. **e,** Sorting the edges in ascending order based on PCC-MR values and dividing them into bins (Methods). **f,** Module identification and gene assignment process. The raw network (bottom layers) is comprised of edges within each bin. Next, the network was filtered using triangles, meaning only edges that form a triangle survive (middle layers): Finally, non-consecutive nodes were processed to obtain the final network (Consecutive network, top layers; see Extended Data Fig.2). Modules are identified using the consecutive network, and genes are assigned to modules using the triangle network. This process is repeated for each bin. **g,** Enlarged examples of the raw network, triangle network, and consecutive network for a single bin. In the consecutive network, clusters with different colors denote identified modules. White nodes represent unassigned genes. These colors are mapped back onto the triangle network, with black nodes representing non-consecutive (eliminated) nodes.

In CGMFinder, a gene’s assigned module is its most highly connected module. For each bin, after the identification of new modules, genes are assigned to modules based on their connections with these identified modules. For a particular gene, all modules connected to the gene are first examined in the raw network. The module with the highest number of connections to this gene is selected as a candidate module, provided its connection count is at least twice that of the second-most connected module. The gene’s association with the candidate module is then further evaluated in the triangle network. The gene is finally assigned to the module if it meets a specific user-defined ratio of connections with members of the candidate module. Genes that do not meet these criteria remain unassigned (Fig. 2g). This process is repeated for each bin until all edges are integrated into the network and all genes are interrogated.

### Validation of modules identified in the MALBAC-DT dataset

To assess the performance of CGMFinder, we initially applied it to our 3H GM12878 dataset, which was generated using the MALBAC-DT method. This dataset includes 4,130 cells that were profiled and deeply sequenced, resulting in a zero count rate of 40%. A total of 63 modules were identified (Fig. 3a), encompassing approximately 3,500 genes assigned to these modules. As expected, more highly expressed genes were assigned to modules than those with lower expression (Extended Data Fig. 3).

**Fig. 3.**
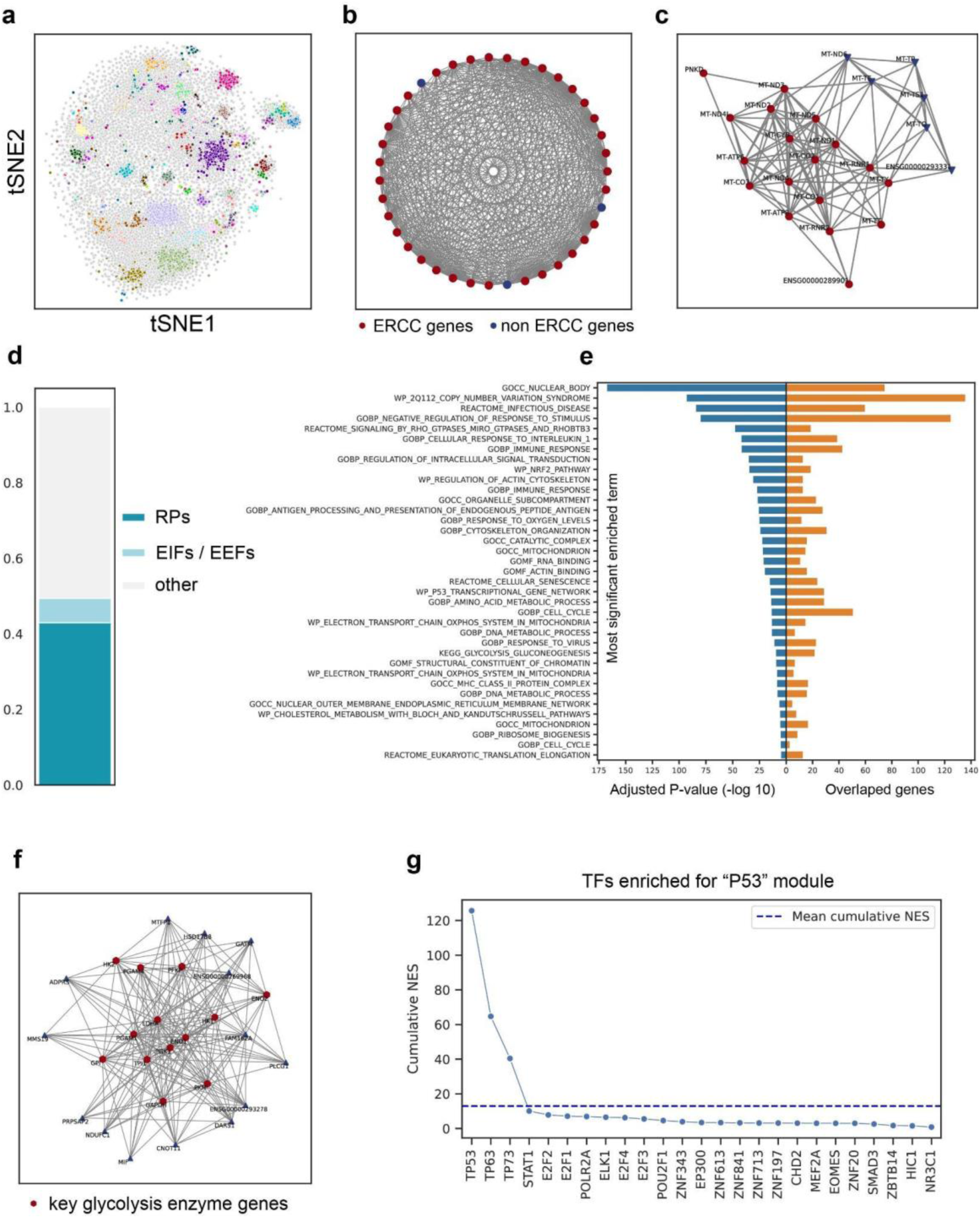
Validation of modules identified in GM12878. **a,** t-SNE plot of identified CGMs. Genes are color-coded to depict modules identified by CGMFinder, with gray dots representing unassigned genes. **b,** Gene composition of the ERCC module, with 36 ERCC genes assigned (red dot) and 3 false positives (blue dot), lines representing connections among nodes in the consecutive network. **c,** Gene composition of the two mitochondria modules identified **d,** Ratio of RPs and Eukaryotic translation Initiation Factors (EIFs) or Eukaryotic translation Elongation Factors (EEFs) in the identified “ribosome module”. **e,** Over Representation Analysis of the top forty enriched modules. The name of the most significantly enriched functional term is shown for each module. Adjusted P-values are shown in the left panel, and the number of overlapping genes is shown in the right panel. **f,** Gene composition of the “glycolysis module”. Red dots represent key enzymes responsible for glycolysis. **g,** The Normalized Enrichment Score (NES) is a Z-score reported by the cisTarget database indicating the degree of motif enrichment. For a specific enriched TF, all motif hits with an NES greater than 3 are summed to calculate the “cumulative NES” (Methods). Members of the TP53 family, which exhibit a cumulative NES significantly above the mean, are specifically enriched within the “p53 transcriptional gene network” module.

We took several approaches to evaluate the accuracy and validity of modules detected by CGMFinder. Initially, the gene composition of the two ground-truth modules was examined. We asked how many of the 92 ERCC spike-ins were assigned to a module in common. Of the 92 ERCC spike-ins, we kept the 39 most abundant RNAs detected in more than 10% of cells for analysis. Of these 39 ERCC spike-ins, 36 were assigned to an ERCC module (True positive) while 3 genome-encoded genes were inaccurately included (False positives), resulting in a detection sensitivity of 0.92 and precision of 0.92 (Fig. 3b). The three ERCC genes not assigned to the ERCC module (False negatives) were left unassigned, effectively controlling potential false positives. Genes encoded by the mitochondrial genome (internal standard) were assigned to two distinct modules: one predominantly comprising heavy strand genes and the other mainly consisting of light strand genes. The distinction between the heavy-strand and light-strand genes makes sense mechanistically since polycistronic genes encoded on opposite strands are transcribed independently of the other at the molecular level, while at the same time likely to exhibit high correlation given their shared mitochondrial function. All 21 mitochondrial genes detected (13 protein-coding, 2 rRNA, and 6 tRNA) are allocated across these two modules, along with 3 genome-encoded genes, resulting in a detection sensitivity of 1 and a precision of 0.875 (Fig. 3c).

We next utilized Over Representation Analysis (ORA) to assign biological functionalities to the modules identified by CGMFinder. Among the identified CGMs, 85% exhibit an adjusted p-value below 0.05, with 72% displaying an even lower adjusted p-value of less than 0.01 (Fig. 3e). These modules encompass a diverse array of functions, including encoding protein complexes, involvement in metabolic pathways, and contribution to specific biological processes. We will showcase a selection of these modules as examples to compare them with established knowledge, validating the accuracy of module identification. One canonical example is the tight co-regulation of genes encoding components of the ribosome, a vital cellular structure responsible for protein synthesis. The CGMFinder identified a “ribosome module” harboring 71 out of the 80 known ribosomal proteins (RPs) genes in human cells42, representing 89% of reported RP genes. Further, seven translation initiation factors (EIF2A, EIF2S3, EIF3E, EIF3F, EIF3H, EIF3K, EIF4B) and four translation elongation factors (EEF1A1, EEF1B2, EEF1G, EEF2) are included in this module (Fig. 3d). Likewise, the glycolysis pathway orchestrates, a fundamental metabolic process for ATP generation from glucose. CGMFinder identified a specific module designated as the “glycolysis module.” This module includes nine out of the ten canonical genes responsible for catalyzing glycolysis reactions43, along with LDHA, which converts excess pyruvate to lactate (Fig. 3f). A third noteworthy example is related to the MHC class I antigen presentation pathway44,45, pivotal in presenting peptide fragments on cell surfaces. The “MHC class I module” comprises genes like HLA-A, HLA-B, HLA-C, HLA-E, and B2M encoding proteins forming the MHC I complex, alongside others like PSMB9, TAP1, and TABP (Tapasin) involved in antigen processing and loading. For a detailed list of all identified modules, please refer to the Extended Data.

Transcription factors (TFs) play a crucial role in orchestrating gene expression within modules. Hence, conducting TF enrichment analysis for CGMs acts as a complementary approach to validate the accuracy of module identification. In this study, the cisTarget database^46–48^ was employed to identify regulatory TFs for CGMs (Methods). For example, the “p53 transcriptional gene network” module, comprising 83 genes with 51 overlaps with p53 direct targets^49^, is primarily enriched for the p53 TF family (Fig. 3g). Likewise, the “response to oxygen levels” module showed primary enrichment for HIF1A and ARNT, crucial in cellular responses to oxygen levels, consistent with prior research^50^ (Fig. 6f). Enriched TFs for the MHC class II CGM included members of the NF-κB family, such as RELA, NFKB1, NFKB2, as well as RFX5 and NFYA, all of which are reported to regulate MHC class II gene expression^51,52^ (Extended Data Fig. 4b). Additionally, the enrichment analysis of the “ribosome biogenesis” module highlighted the regulatory roles of transcription factors MYC and MAX, supported by existing literature ^53–55^ (Extended Data Fig. 4c).

### Deciphering functional organization through module connectivity analysis

Genes do not operate in isolation, and this principle similarly applies to modules with diverse biological functions. Modules are often coordinated to perform essential functions or to adapt to environmental changes. By investigating the connections among these modules, we can attain a comprehensive understanding of the biological system as a whole at higher organizational levels.

In CGMFinder, we introduce a method for measuring the connections between different modules. The “connectivity” between two modules is defined as the ratio of inter-module connected genes to the total number of genes in these two modules in the triangle network (Extended Data Fig. 5a). At each bin, the connectivity of each module to all the other modules is computed repeatedly. This process yields a connectivity curve illustrating how the connectivity changes along the axis of edge bins (Extended Data Fig. 5b). Subsequently, the Area Under Curve (AUC) is calculated for each module, which is then used to generate an AUC matrix (Extended Data Fig. 5c). A higher AUC value signifies a stronger connection between two modules. Furthermore, the AUC matrix is converted into an AUC MR matrix, which serves as a distance matrix (Extended Data Fig. 5d), utilizing a method similar to that used in gene MR matrix calculations. Hierarchical clustering is then applied to group the modules and form a dendrogram, providing valuable insights into their relationships and clustering patterns (Extended Data Fig. 5e)

In the dendrogram, some modules show linkage at the distance value of one. This proximity suggests a mutual top-ranking connection between each other, potentially highlighting related functions or shared characteristics. For example, two modules from the mitochondrial genome—one from the heavy chain and the other from the light chain—exhibit a close connection at this distance, corroborating our expectation of a strong link due to their common origin (Fig. 3c and Fig. 4a-Ⅰ). Similarly, the “glycolysis” module and the “response to oxygen levels” module, both crucial for cellular aerobic metabolism^56^, also display a close relationship (Fig. 4a-Ⅱ). Furthermore, two antigen presentation modules, MHC class I and MHC class II, which are responsible for presenting antigens internally and externally within the cell, also demonstrate a tight association (Fig. 4a-Ⅲ). Additional examples of modules exhibiting similar close associations are also observed (Fig. 4a).

**Fig. 4.**
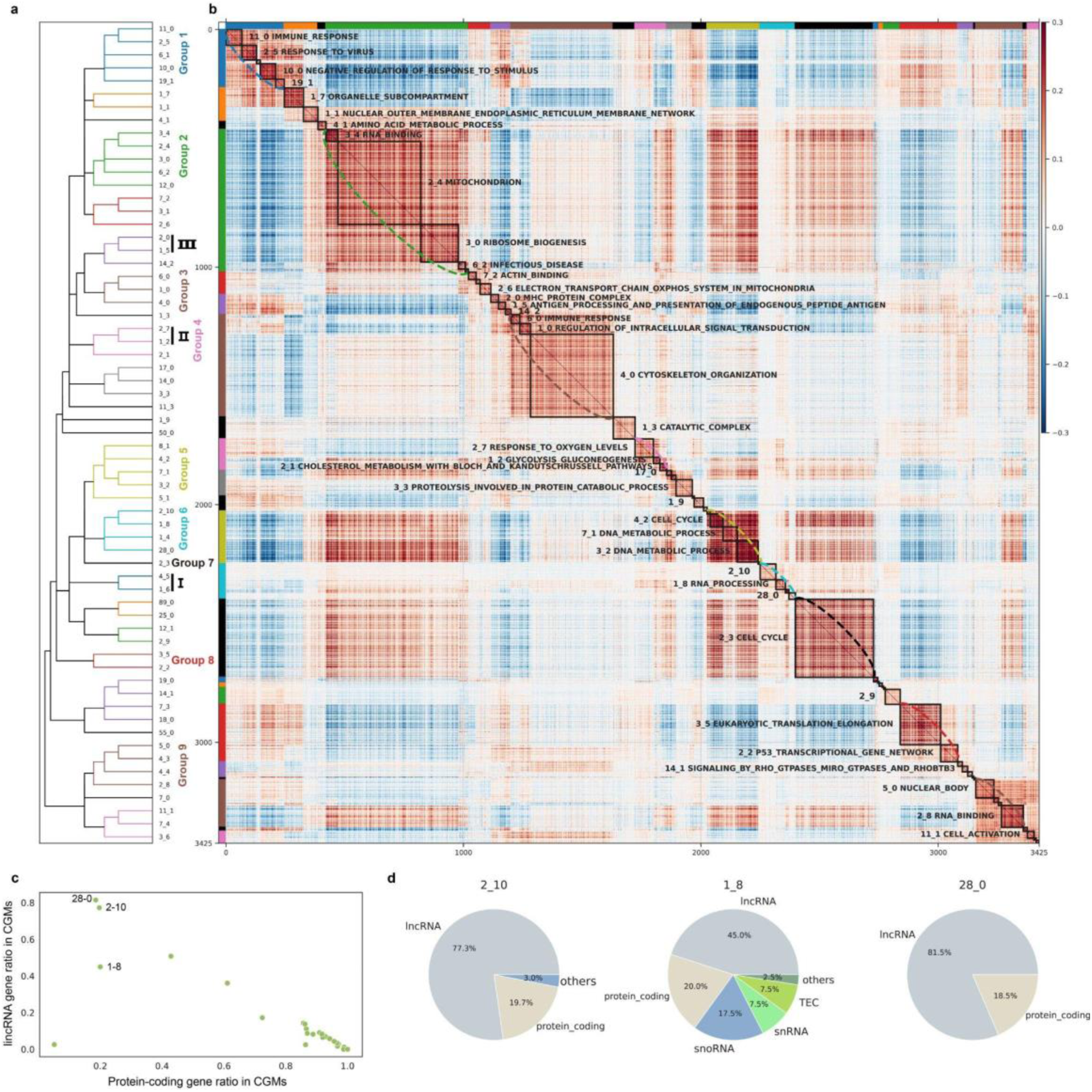
Modular structure in GM12878. a, Hierarchical clustering of CGMs based on module connection analysis. Module groups are color-coded (Y-axis) with the function of selected groups indicated by Roman numerals (see main text). Individual modules are named in the format 4_0, where ‘4’ indicates the bin number and ‘0’ indicates it is the first module within that bin. b, Heat map displaying the Pearson correlation coefficients (PCC) of genes assigned to CGMs. Modules are enclosed in squares along the diagonal, maintaining the same order as in (a). Module groups are indicated by colors along the left side and the top of the plot, and by a dashed arc line along the diagonal. On the graph, the module name is followed by the top significantly enriched functional term. c, Distribution of the proportion of lncRNA in CGMs identified in GM12878. Several modules are comprised mainly of non-coding genes. d, Pie chart displaying the gene composition of three modules in (c) containing a high ratio of lncRNA.

Examination of the dendrogram at a higher distance reveals several module groups, each containing modules with similar biological functions. (Fig. 4a,b). Group two primarily consists of modules related to “housekeeping” functions, including “mitochondrial protein-containing complex,” “ribosome biogenesis,” and “spliceosomal complex” (Fig. 4a,b, group 2). Groups five and seven primarily contain modules associated with cell cycle functions. Group five encompasses modules associated with G1/S “cell cycle” functions, approximately 80% of the G1/S phase marker genes reported^57^ are contained in this group. (Fig. 4a,b, group 5). Group seven, which contains a large single module annotated as “mitotic nuclear division”, focuses on genes related to the G2/M cell cycle phase. Approximately 90% of the G2/M phase marker genes reported^57^ are contained in this module (Fig. 4a,b, group 7).

Group six, which is adjacent to the two cell cycle groups, contains modules that possess a significant proportion of lincRNA genes (Fig. 4a,b, group 6; Fig. 4c,d), which may suggest their important role in cell cycle progression. Groups one and three are characterized by modules with functions specific to the immune response. Group one includes modules related to “immune response regulating signaling pathway”, “negative regulation of viral process” and “autophagosome organization” (Fig. 4a,b, group 1). Group three, which is adjacent to the two antigen presentation modules, includes modules involved in the “B cell activation”, “NF-κB signaling”, and “actin filament-based process” (Fig. 4a,b, group 3).

Group four consists of modules related to “response to oxygen levels”, “ADP metabolic process (glycolysis)” and “sterol biosynthesis process” (Fig. 4a,b, group 4). The grouping of these modules reflects the coordinated regulation of energy production under hypoxic conditions, a pattern not observed in normoxia (discussed later). Group eight contains the “ribosome module” and “p53 module”. Finally, group nine primarily consists of genes from chromosome one, which may be attributed to aneuploidy or copy number variations of chromosome (Fig. 4a,b, group 9; Extended Data Fig. 6). Analyzing module connections provides insights into the functional associations between modules, thereby enhancing our understanding of the biological processes under investigation.

### Effects of cell number, sequencing depth, and aneuploidy on module detection

The number of detected modules and assigned genes is significantly influenced by the number of profiled cells and sequencing depth. Through downsampling analysis, we investigated their effects on module detection.

Generally, increasing the number of profiled cells significantly reduced the number of “noisy modules”, as indicated by a higher ratio of functionally annotated modules (Fig. 5a). This effect is particularly pronounced when the number of profiled cells is below one thousand, as limited statistical power can result in uncorrelated gene pairs being incorrectly measured as correlated (Fig. 1a). In contrast, increasing the sequencing depth per cell significantly enhances the number of identified modules, as more genes are accurately detected with higher sequencing depth. However, greater sequencing depth can also lead to an increase in noisy modules, potentially due to inaccuracies in quantifying low-expressed genes, which are not detected with lower sequencing depth (Fig. 5b). Both increasing the number of cells and improving sequencing depth lead to a continuous rise in the number of genes assigned to modules, as indicated by the increased ratio of assigned genes. (Fig. 5c).

**Fig. 5.**
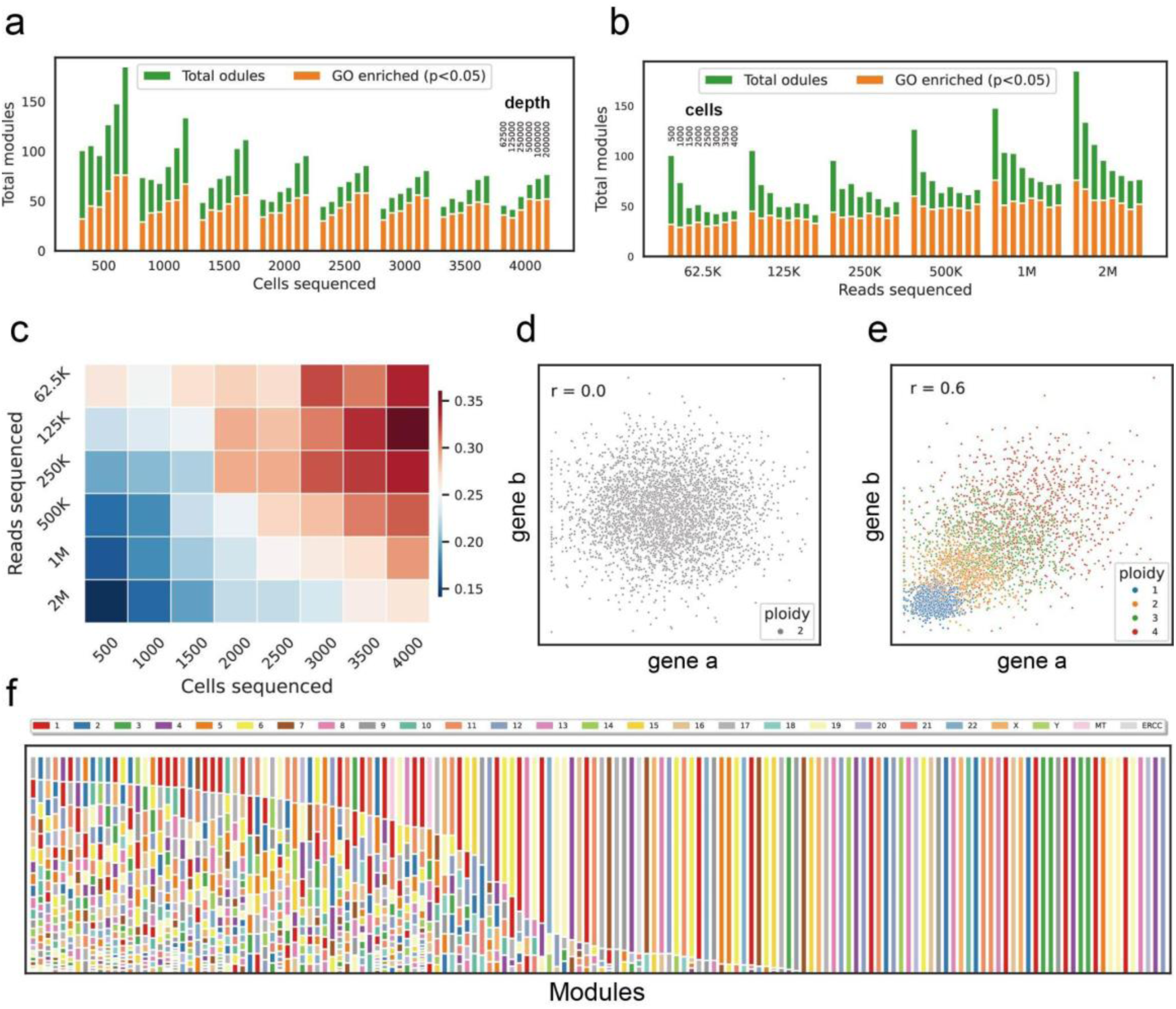
Factors affecting module identification. a, b Each vertical bar represents the number of CGMs detected across the indicated combinations of (a) downsampled cell number, with increasing sequencing depth as indicated or (b) downsampled sequencing depth, with increasing cell numbers as indicated. The orange portion of each bar shows the number of CGMs that contain a functionally enriched GO term. c, Proportion of genes assigned to modules across different combinations of downsampled cells and sequencing depth. d, e Simulation demonstrating the effect of aneuploidy on correlation measurement for genes located on the same chromosome. Two genes located on the same chromosome that would not be correlated in diploid cells (d) could exhibit high correlation in populations with variable ploidy of that chromosome(e). Each simulated cell is assigned a random ploidy of 1 to 4, and the expression of genes a and b are adjusted accordingly. f, Chromosomal sources of genes within modules detected in the K562 dataset. Modules originating from a single chromosome are likely due to a variable and unstable copy number of that chromosome in the cell population.

Another factor influencing module detection is the presence of aneuploidy in cells. Aneuploidy disrupts gene expression levels by altering gene dosage, thereby affecting the correlation measurements of gene pairs. Genes that would otherwise show no correlation may appear correlated due to altered ploidy in different cells (Fig. 5d,e). To examine the impact of aneuploidy on module detection, we generated a dataset for the K562 cell line using MALBAC-DT, which is known for its widespread aneuploidy and various chromosomal structural abnormalities ^58^. In the K562 dataset, we observed a higher number of modules enriched with genes from the same chromosome (Fig. 5f), a phenomenon more pronounced compared to the GM12878 dataset (Extended Data Fig. 6).

### Comparison of modules identified from datasets generated by MALBAC-DT and 10x

To assess the performance of CGMFinder across datasets produced by different scRNA-seq platforms, we compared modules identified from three different datasets for the same GM12878 cell line. The first dataset is the MALBAC-DT dataset, comprised of about 4,000 cells and contains approximately 40% zeros in the gene expression matrix. The second dataset, derived from an earlier study ^59^, was generated using 10x v2 chemistry, comprising about 7,000 cells and containing about 60% zeros in the expression matrix. The third dataset, created in this study using the latest 10x v4 chemistry, includes approximately 10,000 cells with roughly 30% zeros.

As previously discussed, increased sequencing depth and cell numbers enhance module detection and gene assignment. The 10x v4 dataset, which has the highest sequencing depth and cell count, identified the most modules and assigned the largest number of genes. In contrast, the 10x v2 dataset, with the lowest sequencing depth, detected the fewest modules. The MALBAC-DT dataset detected more modules than the 10x v2 dataset but assigned fewer genes. This difference can be attributed to the MALBAC-DT dataset’s greater sequencing depth but lower number of profiled cells (Fig. 6a,b).

Focusing on the ground-truth mitochondria module, the 10x v4 dataset detected the most mitochondria genes with a sensitivity of 0.82 and a precision of 0.93. The 10x v2 dataset detected the fewest mitochondria genes and had a much lower precision of 0.6, despite maintaining high sensitivity (Fig. 6c,d). The MALBAC-DT dataset achieved a sensitivity of 1 and a precision of 0.875 for mitochondria module identification.

Overall, the modules detected in these three datasets are highly comparable. For example, 81% of the modules identified by MALBAC-DT have an Overlap Coefficient greater than 0.2 when compared to modules detected in the 10x v4 dataset. This overlap is 79% for the 10x v2 dataset compared to the 10x v4 dataset, and 72% for the 10x v2 dataset compared to the MALBAC-DT dataset (Fig. 6e). Despite this general comparability, each dataset identifies some specific modules unique to it. The 10x v4 dataset identified 36 and 63 unique modules compared to the MALBAC-DT and 10x v2 datasets, with 64% and 67% of these modules being functionally enriched. The MALBAC-DT dataset found 12 and 34 unique modules compared to the 10x v4 and 10x v2 datasets, with 42% and 53% of these modules being functionally enriched. The 10x v2 dataset also identified 12 and 9 unique modules compared to the 10x v4 and MALBAC-DT datasets, with 75% and 78% of these modules being functionally enriched.

**Fig. 6.**
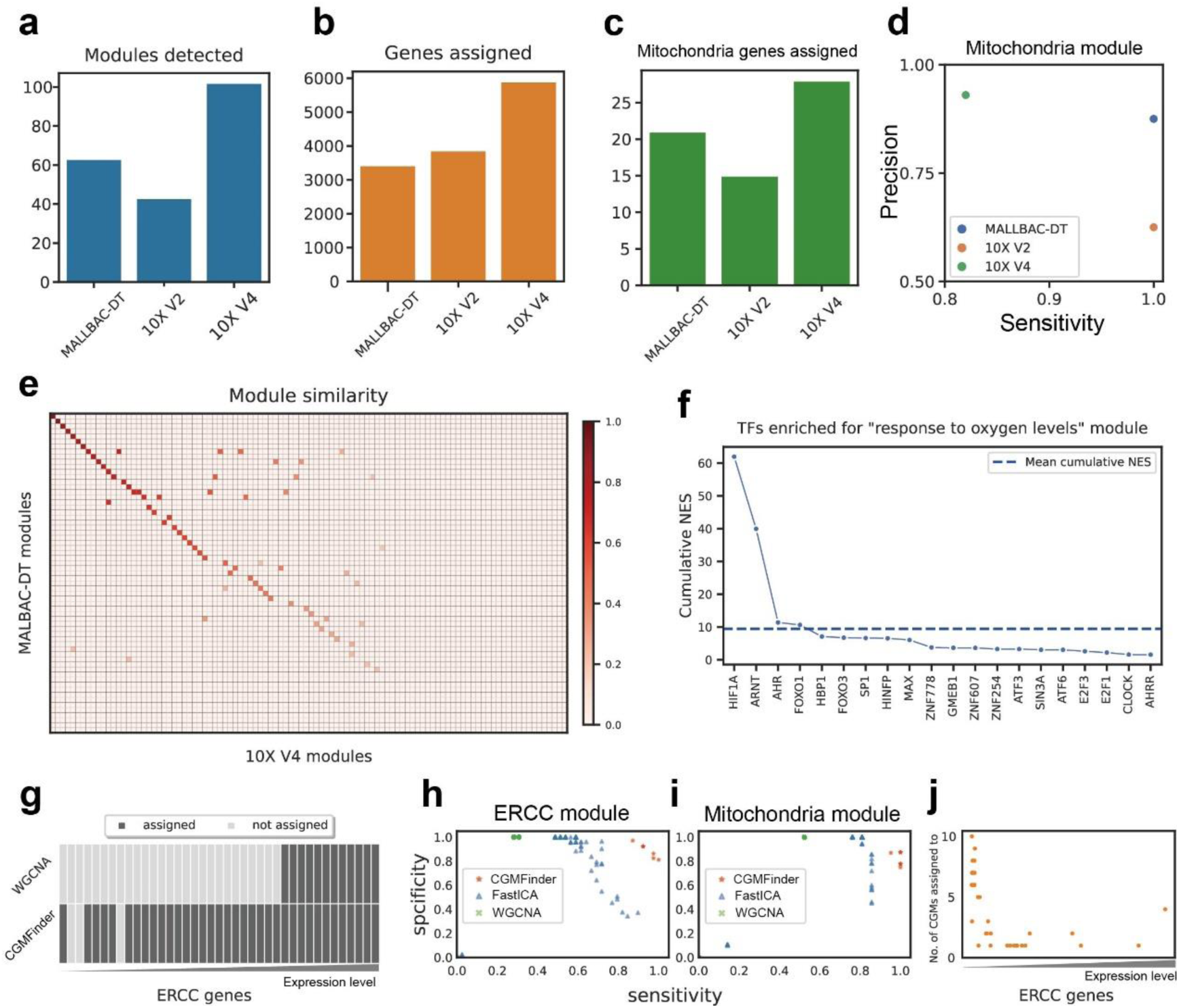
Performance comparison for different datasets and module identification methods. a,b,c Comparison of modules identified with the in-lab MALLBAC-DT protocol, and the commercial 10X Genomics version 2 and version 4 kits. (a) Modules detected. (b) Genes assigned and (c) mitochondria genes detected.. d, Sensitivity, and specificity of mitochondria module identification for the three different datasets. e, Similarity (Overlap Coefficient) between modules identified in MALBAC-DT dataset and 10x v4 dataset. f, Enrichment of hypoxia-related transcription factors in the “response to oxygen levels” module. d, ERCC genes assigned by CGMFinder and WGCNA; low-expressed ERCC genes are not assigned in WGCNA. h, i Sensitivity and specificity for ERCC module identification (h) and mitochondria module identification (i) across three different module detection methods. A parameter grid search was conducted for each method to optimize performance. j, Allocation of low-expressed ERCC genes to more modules by FastICA.

Interestingly, two modules functionally annotated as “glycolysis” and “response to oxygen levels,” are uniquely identified in the MALBAC-DT dataset. This may be attributed to hypoxic stress experienced during the prolonged sample preparation and cell sorting processes, regulated by hypoxia-inducible factors (HIFs)^60^. Notably, the two HIFs, HIF1A, and ARNT, are specifically enriched for this module (Fig. 6f). In contrast, the expression of genes in these two modules is not synchronized across the other two datasets. To confirm the presence of these hypoxia-specific CGMs, we treated GM12878 cells under hypoxic conditions and generated another dataset using the 10x platform. Indeed, the two hypoxic-specific modules were reproduced in this dataset after applying CGMFinder for module detection. Additionally, we observed that energy metabolism-related CGMs (such as glycolysis, cholesterol biosynthesis, and mitochondria) cluster distinctly in hypoxic datasets compared to normoxic datasets (Extended Data Fig. 7). This suggests a coordinated regulation of these genes to adapt to hypoxic conditions (Extended Data Fig. 7). The unique modules identified by each dataset may reflect specific physiological variations for each sample.

### Comparison of CGMFinder and other module detection methods

Among the various methods for detecting CGMs, WGCNA stands out as a widely used approach. Initially developed for module detection in bulk RNA-seq data, WGCNA has recently been applied to scRNA-seq data. Additionally, the FastICA algorithm^66^, an Independent Component Analysis (ICA)-based decomposition method, has shown superior performance in gene expression data for module detection through an evaluation of module detection techniques^29^. In our assessment of CGMFinder’s effectiveness, we compare its performance with these two methods. Our evaluation focuses on the detection accuracy of the two ground-truth modules for assessing the performance of each method. The MALBAC-DT dataset was used for comparison (ERCC cannot be spiked in for 10x platform).

For each method, various parameters were tested, and the optimal results were chosen for comparison. CGMFinder achieved a sensitivity of 0.92 and a precision of 0.92 for the ERCC module (Fig 6h). WGCNA showed a remarkable precision of 1 but a lower sensitivity of 0.44; low-expressed ERCC genes were not assigned (Fig 6g,h). The FastICA algorithm achieved the highest sensitivity of 0.76 and precision of 0.88. FastICA allows the assignment of a gene to multiple modules, and low-expressed ERCC genes tended to be assigned to many more modules (Fig 6g,j). It’s worth noting that only CGMFinder achieved both high sensitivity and precision simultaneously. For the mitochondria module, CGMFinder also demonstrated superior performance (Fig. 6i).

## Discussion

Identifying CGMs allows researchers to gain deeper insights into how genes work together and their roles in specific biological contexts. The advent of single-cell RNA sequencing (scRNA-seq) technology has enabled us to explore CGMs at the single-cell level. However, the high sparsity of scRNA-seq data poses significant challenges for accurate module identification. To tackle this issue, we generated a scRNA-seq dataset with the 3H criteria using MALBAC-DT for module identification, which exhibits reduced sparsity and allows for more precise measurement of gene correlations. Additionally, we proposed to use the ERCC module and mitochondria module as ground-truth references to help the selection of correlation measurements and evaluation of the accuracy for identified modules. Based on these, we explored the characteristics of correlation bias, which ultimately led to the development of CGMFinder.

CGMFinder partitions the network into bins and uses triangles to filter the network for robust connections. In this framework, robust highly expressed genes are prioritized for earlier assignment, whereas noisy lowly expressed genes undergo more rigorous interrogation before being assigned. In contrast, WGCNA typically leaves lowly expressed genes unassigned, and FastICA tends to allocate them into a larger number of modules. This is precisely why CGMFinder excels, achieving superior sensitivity and precision in module identification. Evaluation based on the two ground-truth modules indicates that CGMFinder can achieve both high sensitivity and precision. Further, we show that most modules identified in the human B lymphocyte cell line are with specific biological functions. A feature of CGMFinder is its capability to facilitate module connectivity analysis. This aspect enables scCGMTiT to offer insights into the intricate links between modules. Our study revealed groups of modules associated with distinct cellular functions, such as cell cycle regulation, housekeeping processes, and cell type-specific functions. These findings underscore the critical role of module connectivity analysis in unraveling the complex regulatory mechanisms that govern cellular behavior.

Furthermore, we examine the beneficial effects of increasing cell number and sequencing depth on module detection, which reinforces the importance of 3H datasets in this context. Additionally, aneuploidy poses a specific challenge for module detection, which should be considered when working with such samples.

Producing datasets that meet the 3H criteria necessitates both high sensitivity and high-throughput single-cell RNA sequencing (scRNA-seq) methods. Utilizing the 10x platform, a large number of cells can be efficiently profiled economically. Recently, 10x Genomics introduced v4 chemistry, which enhances both throughput and sensitivity. Comparative analysis of modules identified in the MALBAC-DT dataset and the 10x v4 datasets yielded comparable results. Additionally, we successfully replicated the identification of hypoxia-specific modules in cells subjected to hypoxic stress using the 10x platform, confirming the findings in the MALBAC-DT dataset.

## Methods

### Sequencing data generation

#### Cell culture and processing

In this study, a single-cell clone of the GM12878 cell line was generated to create a homogeneous, steady-state sample. Single cells were pipetted into wells of a 96-well plate containing 150 μl of conditioned media, prepared according to the protocol available at “https://genome.ucsc.edu/ENCODE/protocols/cell/human/GM12878-XiMat_Myers_protocol.pdf”. The cells were cultured until confluent, then transferred to an 8-well plate for further expansion, and finally moved to a T25 flask for large-scale culture. The cells were maintained in RPMI 1640 medium supplemented with 2 mM L-glutamine, 15% FBS, and 1% Penicillin-Streptomycin. Before sorting cells by flow cytometry, centrifuge them, wash them twice with DPBS, and filter them through a 40 µm cell strainer. After sorting the single cells into a 96-well plate containing MALBAC-DT lysis buffer, centrifuge the plates and store them at -80°C until amplification. For the 10X experiment, cells were processed according to the manufacturer’s instructions.

#### MALBAC-DT workflow

MALBAC-DT was performed as previously described^15,61^. Initially, 2 µL of lysis buffer containing the RT primer was distributed into 96-well plates, and a single cell was sorted into each well. The plates were then incubated at 72°C for 3 minutes and held at 4°C for RT primer annealing. Subsequently, 2 µL of RT mix was added to each well and incubated at 55°C for 10 minutes to synthesize the first-strand cDNA. After reverse transcription, 2ul exonuclease mix was added and the plates were incubated at 37°C for 30 minutes to digest excess RT primers, followed by incubation at 80°C for 20 minutes to inactive exonuclease I. Next, 24 µL of PCR master mix was distributed into each well for cDNA amplification. Finally, 2 µL of 10 µM Tru2-G-RT primer was added, and an additional 5 PCR cycles were performed. For sequencing library preparation, 5 µL from each well of a plate was pooled, and 50 µL was cleaned up using 0.8x Ampure Beads. Finally, 50 ng of DNA was used for library preparation using the Vazyme TruePrep DNA Library Prep Kit and sequenced on a NovaSeq6000 with a custom read 2 primer.

#### 10x data generation

Single-cell capture and library preparation were performed using the latest Chromium GEM-X Single Cell 3’ Kit v4, following the manufacturer’s detailed protocols. GM12878 cells were loaded onto the GEM-X Chip, aiming for a recovery of 10,000 cells. The cDNA was initially amplified through 10 PCR cycles, after which 500 ng of the amplified cDNA was employed for library preparation. The resulting library was size-selected to target fragments of approximately 400 bp and subsequently sequenced on a NovaSeq 6000 system. For the generation of the hypoxic-stressed library, 5 million cells were transferred into a 1.5 mL LoBind EP tube containing 1.2 mL of culture media. The tube was sealed with parafilm and incubated at room temperature for 4 hours before being loaded onto the chip.

### Data analysis

#### MALBAC-DT data preprocessing

Cell barcodes and UMIs were extracted from the raw FASTQ files and appended to the header lines to create demultiplexed files. The demultiplexed FASTQ files were then mapped to the human genome (GRCh38.p13) using STAR^62^, with the options “– clip3pAdapterSeq AGATCGGAAGAGCAGATCGGAAGAGC” to remove adapter sequences and “–outFilterMultimapNmax 1” to filter out reads that mapped to multiple locations. Reads were assigned to specific genes using featureCounts^63^ with default parameters. Before UMI counting, reads with fewer than 8 thymine (T) nucleotides in the polyT region of the RT primer were removed. UMIs for each gene in a cell were collapsed if they were within a Hamming distance of less than 4. UMIs were counted for each gene across all cells, and the data were merged to generate the gene expression matrix. Finally, cells were filtered based on the following criteria using Scanpy^64^: (1) reads between 0.8 million and 4 million; (2) detected genes greater than 4,000; (3) detected UMIs between 20,000 and 150,000; (4) fraction of primer A greater than 0.8; (5) reads mapping ratio greater than 0.6; (6) fraction of mitochondrial reads less than 0.0125.

#### 10X data preprocessing

For data generated on the 10x platform, raw FASTQ files were processed using Cell Ranger v8.0.1 with default parameters to obtain the raw gene expression matrix. Cells were subsequently filtered based on the distribution of genes detected, UMIs detected, and mitochondrial reads detected using Scanpy.

#### CGMFinder pipeline

CGMFinder was implemented in Python with the gene expression matrix as its only required input. The process consists of several steps: (1) the gene expression matrix is normalized to account for sequencing depth differences and log-transformed using Scanpy, followed by the removal of genes expressed in less than 10 percent of all cells. (2) Pairwise Pearson Correlation Coefficients (PCC) are then calculated for each gene to generate a PCC matrix, with diagonal values set to minus one. (3) Transform the PCC matrix to the PCC-MR matrix. (4) For each gene, connections to the top N (default 50) nearest connected gene neighbors are extracted to construct the network, with PCC-MR values serving as edge weights. (5) All edges are sorted in ascending order by weight and divided into bins; if edges with the same weight do not exceed size M (default 300), edges with the next weight are added until the edge number exceeds M. (6) A graph object is built using graph-tool, and edges from each bin are added cumulatively to reconstruct the raw network, which is then filtered to a triangle network and further to a consecutive network. New modules are identified in the consecutive network as the first occurrence of independent clusters. Genes are assigned to modules based on their connections to modules in the triangle network; for a specific gene, the top-connected module with at least twice the connections of the second most connected module is recognized as the candidate module, and the gene is assigned to this candidate module if it connects to at least a specified ratio (default 0.3) of module members. Module connections are calculated for each module in the triangle network, and this process is repeated for each edge bin.

#### ORA analysis

ORA was conducted using the offline mode of the GSEApy^65^ Python package to determine the enrichment of gene sets from MSigDB (Molecular Signatures Database) collections. Specifically, the analysis utilized gene sets from two collections: C2 - Canonical Pathways and C5 - Gene Ontology (GO).

#### WGCNA

WGCNA was conducted using the WGCNA R package, adhering closely to the guidance provided in its tutorials. The soft-thresholding power was set to 6, determined using the “pickSoftThreshold” function. Module identification was performed using the “blockwiseModules” function, with Pearson correlation (corType = “pearson”) as the basis for gene-gene co-expression. To optimize results for comparison, various parameter combinations (detectCutHeight, deepsplit, and mergeCutHeight) were explored during the tree-cutting process.

#### FastICA

FastICA was performed using the “ica_fdr” function from the GitHub repository hosted by the Saeys Lab, which is available at www.github.com/saeyslab/moduledetection-evaluation.

#### Transcription factor enrichment

Candidate TFs for CGMs were enriched by using the “mc9nr” gene-based cisTarget database. The “gene vs motif” ranking matrix and “motif2tf” matrix were obtained from the Aerts Lab resource portal at https://resources.aertslab.org/cistarget/. The top 3% of ranked genes were selected for the calculation of AUC for CGMs. Motifs with an NES score greater than 3 were mapped to TFs using the “motif2tf” matrix. Only TFs that were directly annotated were retained. For each module, the NES values for the same TF were summed to rank enrichment. TFs that were not expressed in the GM12878 scRNA-seq data were removed.

## Data availability

The raw sequencing data and cleaned expression matrices generated in this study will be publicly available upon publication.

## Code availability

Scripts and detailed instructions for using the method will be available as Python scripts or Jupyter Notebooks upon publication.

---

**Extended Data Fig 1.**
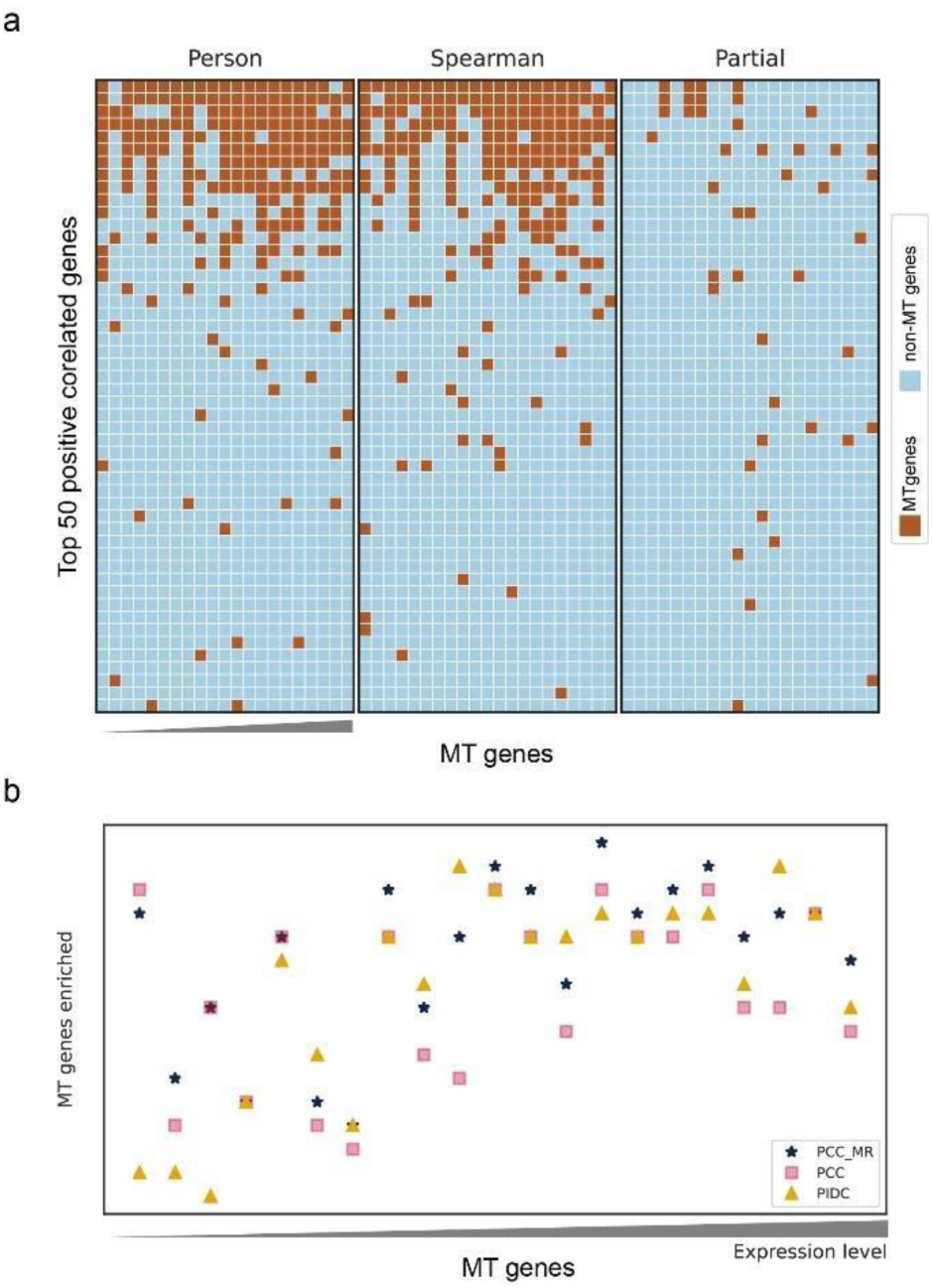
Selection of correlation measurement. a, Top 50 neighbors for the mitochondria (MT) module using PCC, SPC, and partial correlation respectively. Brown dots indicate MT genes and light blue dots represent non-MT genes. b, Enrichment of MT genes in the top 50 positive correlated neighbors for each MT gene using PCC-MR, PCC, and PIDC as measurements. MT genes are sorted by increasing expression levels.

**Extended Data Fig 2.**
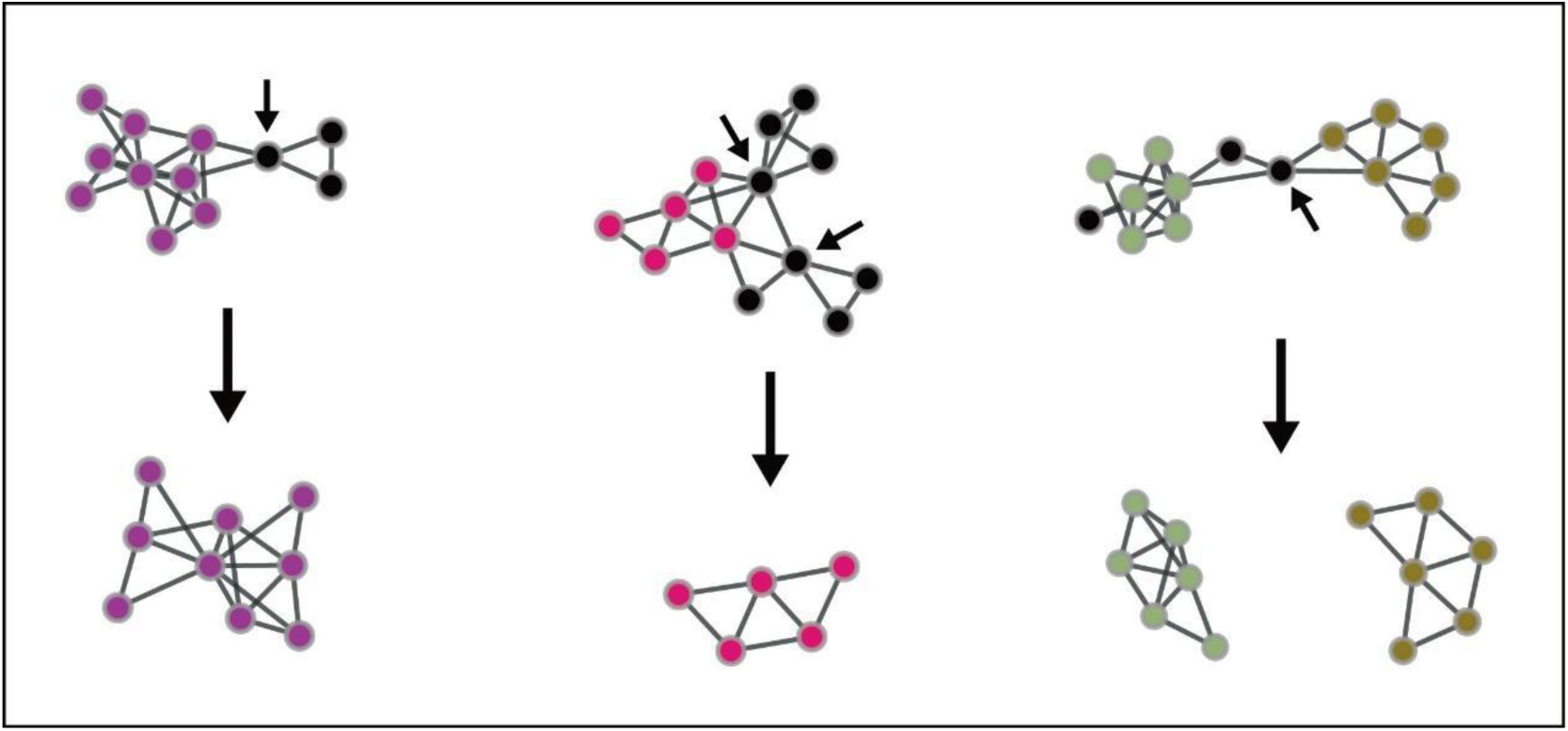
Removal of non-consecutive nodes in the triangle network to produce a consecutive network. Non-consecutive nodes, depicted by arrows in the upper panel, are those whose removal causes clusters to split into at least two parts. After eliminating non-consecutive nodes, the network is re-filtered using triangles.

**Extended Data Fig 3.**
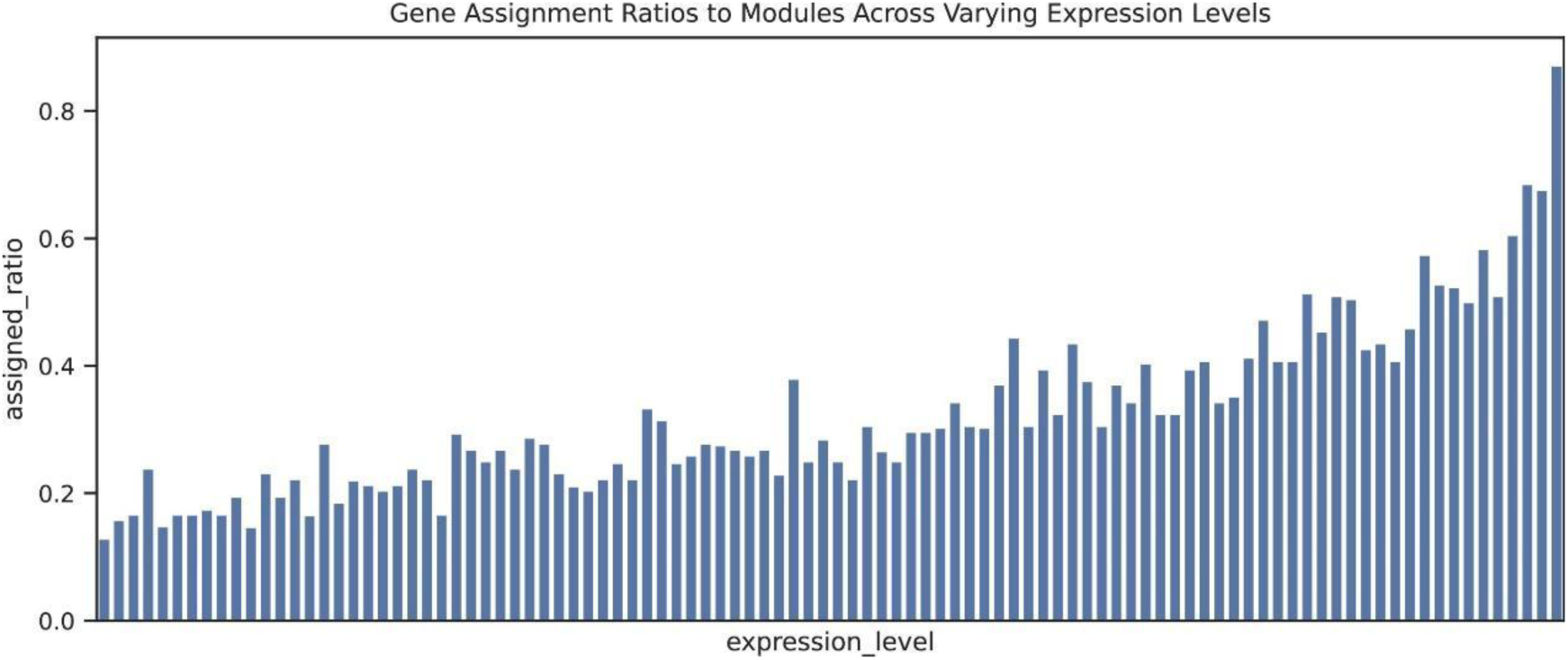
Ratio of genes assigned to modules at different expression levels. Genes are binned according to their expression levels in ascending order. High-expressed genes, which are typically less noisy, exhibit a higher assignment ratio.

**Extended Data Fig 4.**
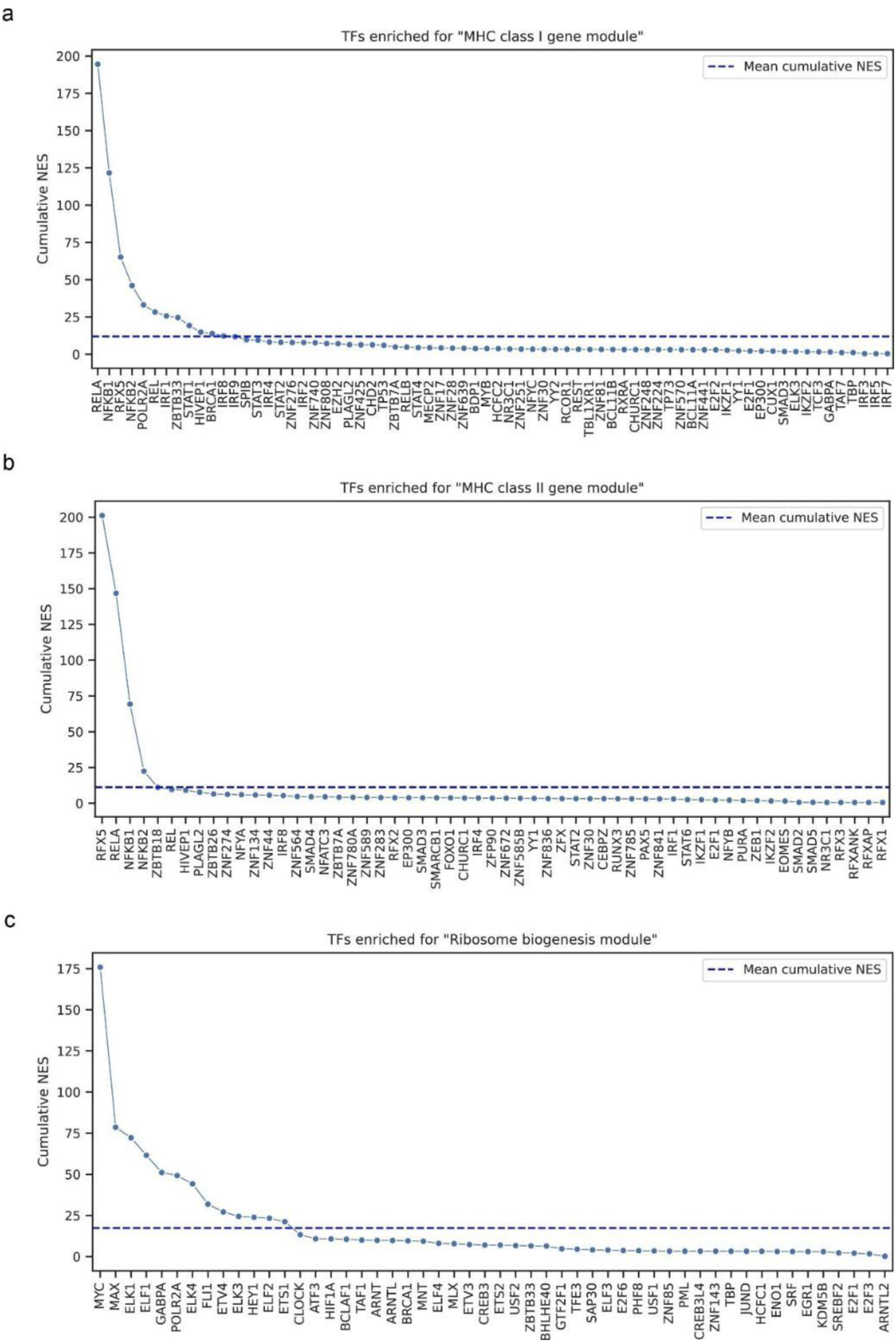
Transcription factors enriched for the “MHC class I module” (a), “MHC class II module” (b), and “Ribosome biogenesis module” (c).

**Extended Data Fig 5.**
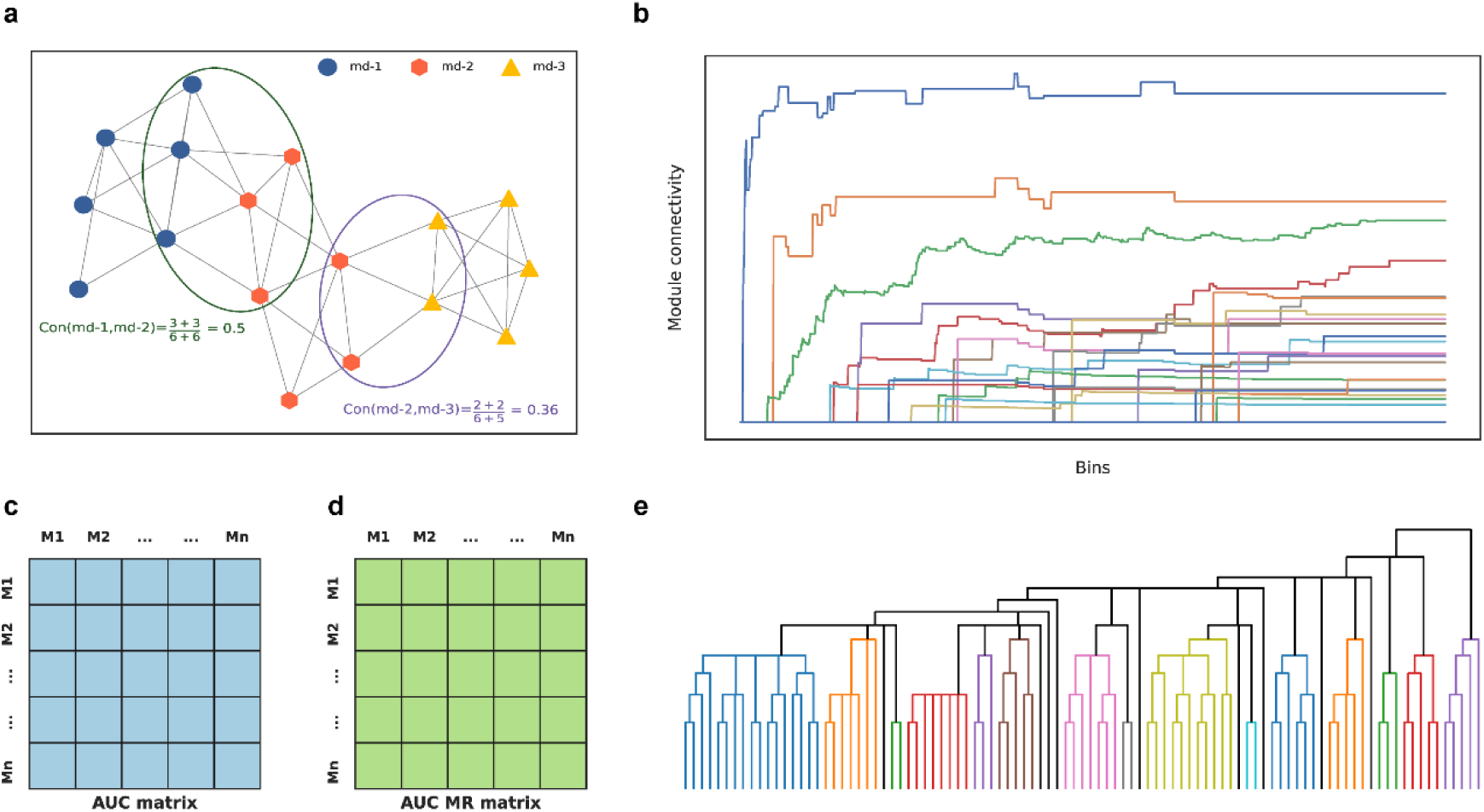
Principle and steps for module connection analysis. a, Definition of connectivity between two modules. b, An example of AUC curve of connectivity for a specific module. The connectivity of this module and all other modules are calculated for each bin and the AUC is calculated for each curve. c, The module AUC matrix is calculated from AUC curve for each module. d, The AUC MR matrix transformed from AUC matrix. e, Hierarchical clustering for modules using module MR values as distance measurements.

**Extended Data Fig 6.**
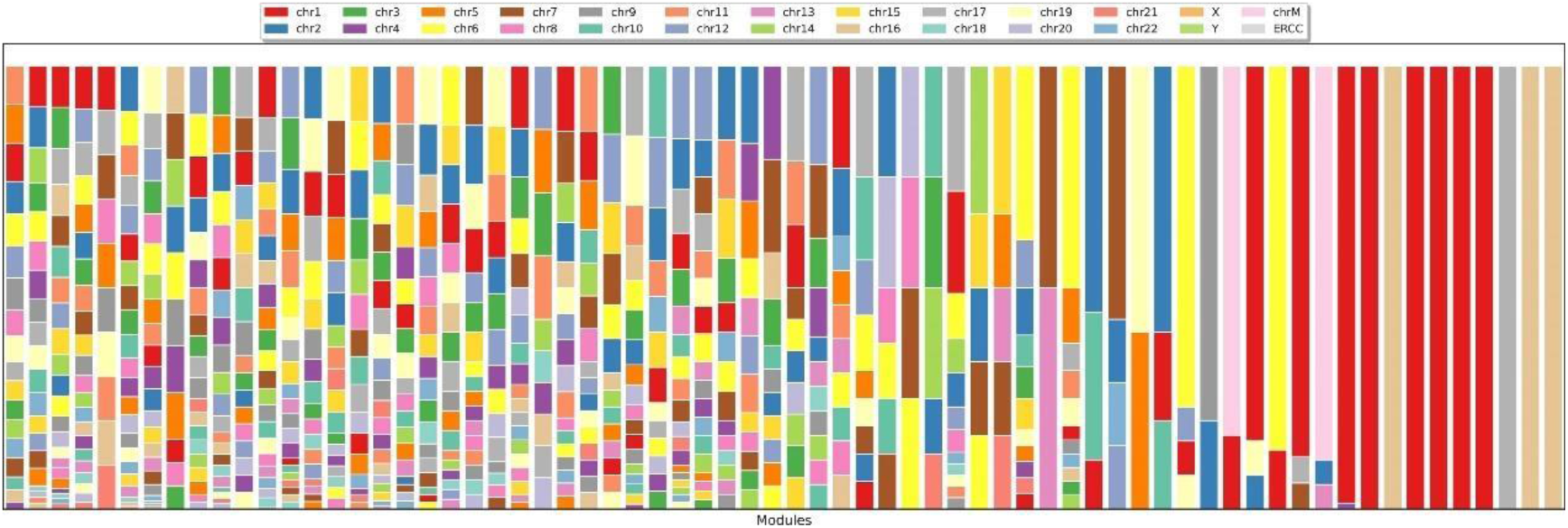
Chromosomal sources of genes within modules detected in the GM12878 dataset. Most modules originate from multiple chromosome sources.

**Extended Data Fig 7.**
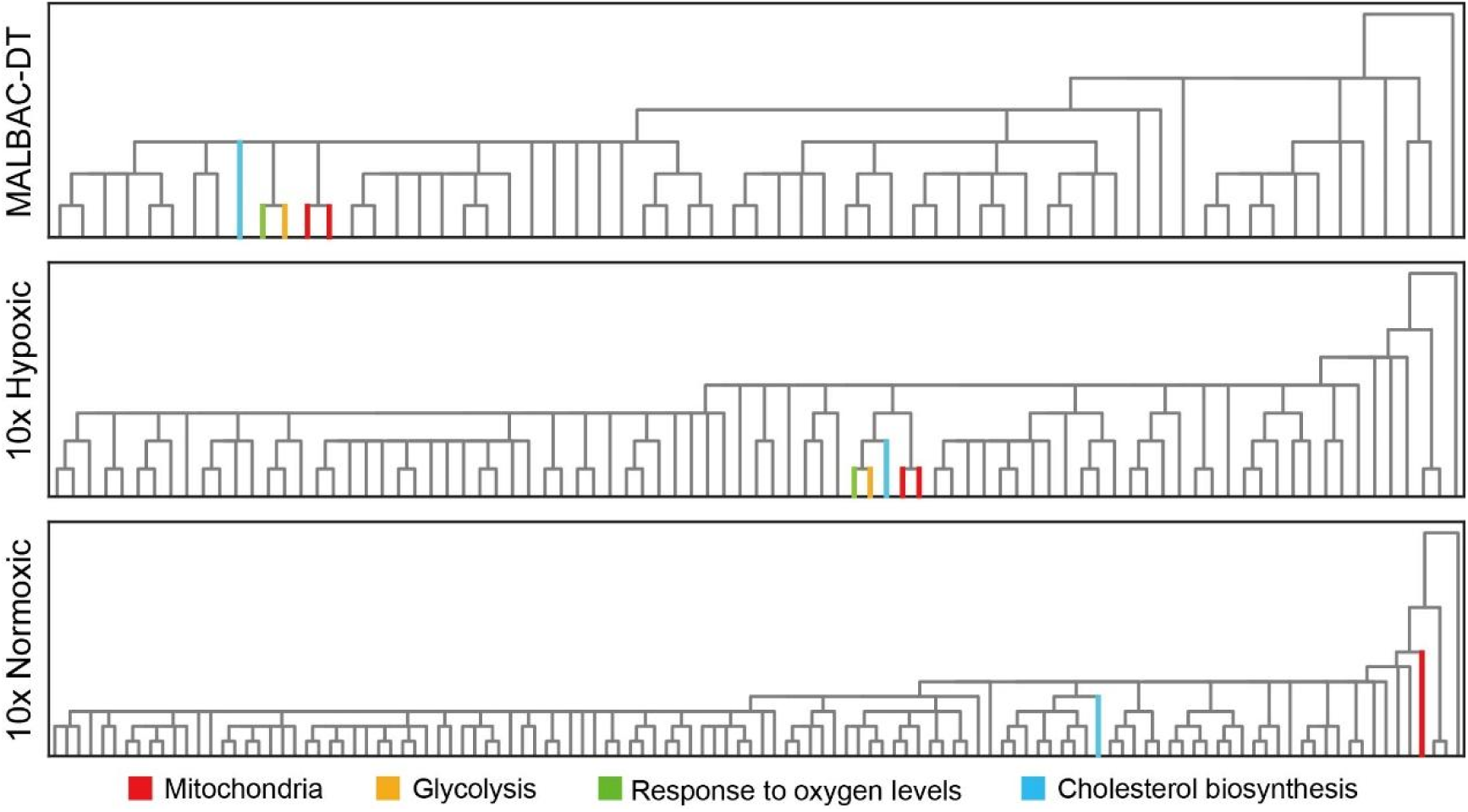
Clustering of energy metabolism-related CGMs in hypoxic datasets. In hypoxic datasets (MALBAC-DT and 10x hypoxic), we observed a distinct clustering of energy metabolism-related CGMs. Notably, the glycolysis module and the “response to oxygen level” module are exclusively present in hypoxic samples. Conversely, in the 10x normoxic sample, the “cholesterol biosynthesis” module and the mitochondria module are significantly separated from each other.

